# Creating microbiome-model harmony between metaproteomics data and the ADM1da for a two-step anaerobic digester

**DOI:** 10.1101/2024.12.20.629579

**Authors:** Patrick Hellwig, Ingolf Seick, Nicole Meinusch, Dirk Benndorf, Jürgen Wiese, Udo Reichl, Robert Heyer

## Abstract

The effective operation, planning, and optimization of renewable energy production in anaerobic digestion (AD) plants relies on advanced process models, such as the Anaerobic Digestion Model No. 1 (ADM1). This study applies an ADM1-based model (ADM1da) to simulate a two-step digester in an industrial setting. The data demonstrate that 2.6% of the methane is lost as a result of open hydrolysis. Conversely, the incorporation of a hydrolysis fermenter enhances methane production by an average of 2.5%. Although ADM1-like models are widely recognized for accurately representing anaerobic digestion processes, mechanistic insights into the microbiome involved have been limited by the absence of tools to analyze microbial composition and functionality at the time these models were developed.

To overcome this limitation, we utilized a metaproteomics approach to assess the abundance and biomass-correlated activity of microbial groups as defined by the model, aiming to bridge the gap between microbial ecology and bioprocess engineering in AD systems. We also developed and evaluated a series of rules for associating particular microbial species with functional groups of the model.

Our analysis demonstrates that while the model supports the presence of a stable microbiome composition in the main fermenter, it is difficult to capture the dynamic behavior observed in the hydrolysis fermenter. Furthermore, the actual AD microbiome displays a greater versatility than the model assumes, with microorganisms performing multiple functions rather than being restricted to single roles.

In conclusion, this study identifies options for improving AD models and integrating comprehensive biological knowledge to further optimize the performance of anaerobic digesters.

**Highlights:**

- Simulations revealed 2.6 % methane volume loss attributed to open hydrolysis
- Implementation of a two-step process increased methane production by 2.5%
- Identification of rules to map metaproteomics data to ADM1da
- Simulations of ADM1da depict the dynamic in the main but not in the hydrolysis fermenter
- Microbial species perform multiple functions not just one as assumed in the ADM1da

## 1. Introduction

The conversion of waste to biogas by complex microbiomes in anaerobic digestion is a key technology for renewable energy production. The microbiomes in anaerobic digesters (AD) microbiomes consist of about 10-20 high-abundant (above 1% of the microbial biomass/cell count) and partly redundant archaeal or bacterial key species and a multiple of further low abundant species (Hassa et al., 2023; Heyer et al., 2024a).

These microbiomes degrade the biomass in the four main steps hydrolysis, acidogenesis, acetogenesis, and methanogenesis. The complexity of the microbiomes and the AD process is further complicated by two challenges. (i.) The microbiome and the AD process depend on parameters such as the feedstock, pH values, temperature, and AD design (Heyer et al., 2016; Vrieze et al., 2015). (ii.) Within the microbiome, plenty of interactions between the different microbial species including syntropy, competition, and phage-host interactions exist (Heyer et al., 2019). To keep an overview of the AD process and to have a powerful tool for operating, planning, and optimizing AD, reduced, sophisticated process models such as the Anaerobic digestion model 1 (ADM1) (Batstone et al., 2002) and its extensions such as ADM1da (Karlsson, 2017; Ogurek et al., 2013) were developed.

The ADM1 is a mathematical model describing the physicochemical and biochemical processes (disintegration, hydrolysis, acidogenesis, acetogenesis, and methanogenesis) of the anaerobic digestion process and seven functional microbial groups (Figure 1). Therefore, the ADM1 comprises ordinary differential equations (ODEs) and differential algebraic equations, covering 26 dynamic state concentration variables, and 8 implicit algebraic variables per reactor vessel (Batstone et al., 2002).

**Figure 1:**
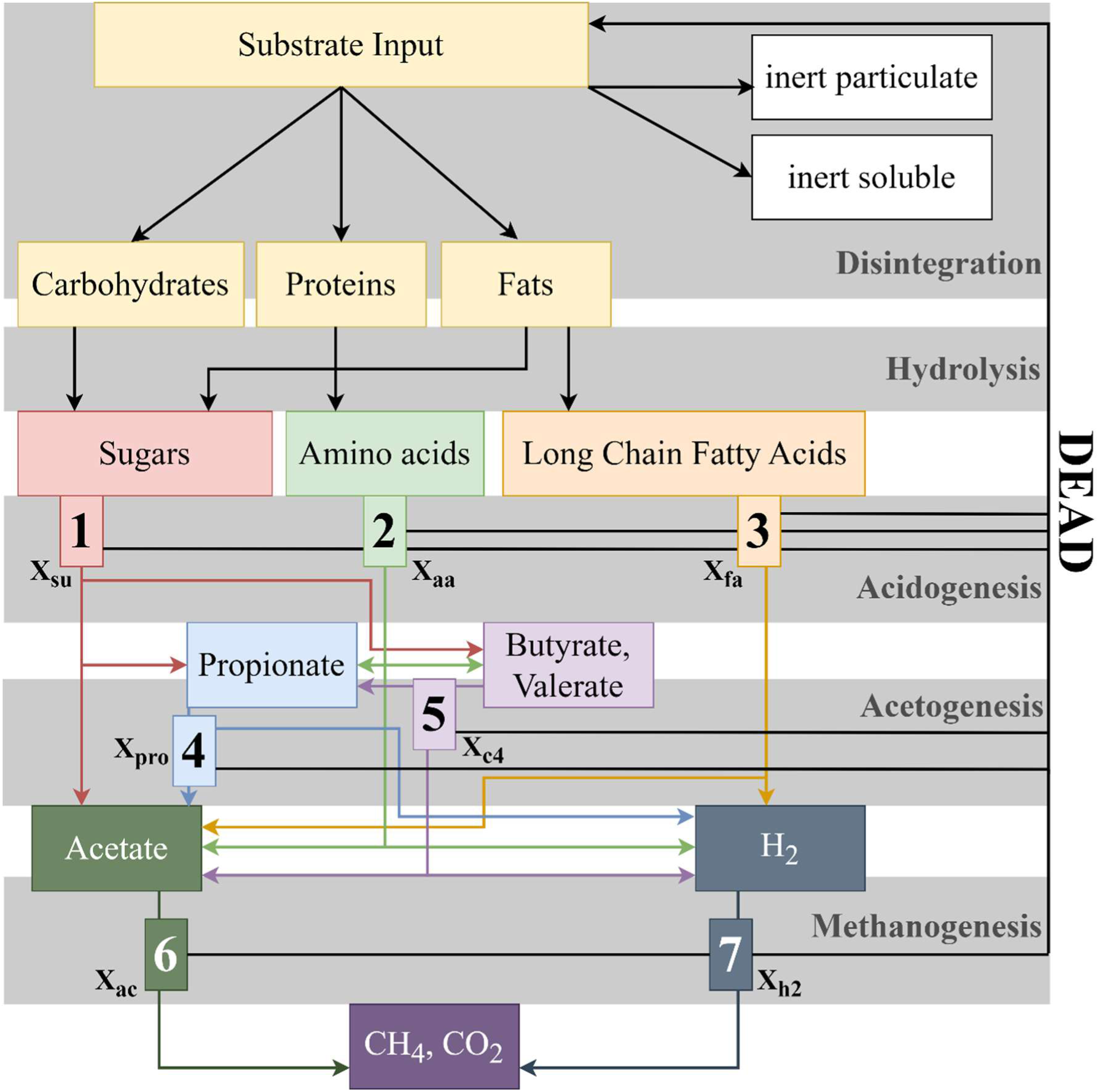
Biochemical processes of the ADM1da. (1) acidogenesis from sugars, (2) acidogenesis from amino acids, (3) acetogenesis from long chain fatty acids, (4) acetogenesis from propionate, (5) acetogenesis from butyrate and valerate, (6) acetoclastic methanogenesis, and (7) hydrogenotrophic methanogenesis. Modified from Karlsson (2017) and Batstone et al. (2002). x_su_: sugar degraders, x_aa_: amino acid degraders, x_fa_: fatty acid degraders, x_c4_: valerate and butyrate degraders, x_pro_: propionate degraders, x_ac_: acetoclastic methanogens, x_h2_: hydrogenotrophic methanogens.

Based on the ADM1 and its adaptations (Mo et al., 2023), researchers investigated different fermenter settings (Bensmann et al., 2013; Blumensaat and Keller, 2005), dynamic processes (Bensmann et al., 2016), the potential of biological methanation (Bensmann et al., 2014), increased AD plant efficiency or integration of the biogas process to complex biomass cycles (Meinusch et al., 2021; Seick et al., 2023). Despite all the previous research studies (e.g., 272 publications found under the term ”ADM1” on Pubmed, 18 September 2024), there are still weaknesses of the model requiring further improvement. For example, hydrolysis is assumed to be a rate-limiting step, but it is debated if it should be described by a simple first-order kinetic or by the inclusion of the biomass of hydrolyzing microorganisms using a Contois kinetic (Contois, 1959; Wang and Li, 2014). One reason for these uncertainties was caused by the inability to analyze the underlying microbiomes in detail. So far the abundance of microorganisms within the ADM1da based on the chemical oxygen demand (COD) is considered by assuming seven groups of microorganisms conducting either (i.) sugar degradation (x_su,_ acidogenesis from sugars), (ii.) amino acid degradation (x_aa_, acidogenesis from amino acids), (iii.) long chain fatty acid degradation (x_fa_, acetogenesis from long chain fatty acids), (iv.) valerate and butyrate degradation (x_c4_, acetogenesis from butyrate and valerate), (v.) propionate degradation (x_pro_, acetogenesis from propionate), (vi.) acetate degradation (x_ac_, acetoclastic methanogenesis), and (vii.) hydrogen degradation (x_h2_, hydrogenotrophic methanogenesis) (Figure 1).

Thanks to the development of novel omics methods, nowadays the taxonomic profile and function of microbiomes using metagenomics (Campanaro et al., 2020) and metaproteomics (Heyer et al., 2015) can be analyzed and could be linked with the ADM1da model (Heyer et al., 2019). Whereas metagenomics has a better resolution, enables the prediction of metagenome-assembled genomes (MAGs) (Bowers et al., 2017), and correlates better with the cell number, metaproteomics correlates better with biomass and metabolic function enabling to estimate of the abundance of microbial metabolic groups assumed by the ADM1da model. Recent metagenomics and metaproteomics data from a two-step industrial-scale AD system (Heyer et al., 2024a) allow for the alignment of the actual biomass composition and dynamics with those predicted by the ADM1da model over a one-year operational period. Building on this dataset, our study aimed to identify strategies for linking microbiome data to the ADM1da model, with the goal of providing new insights to better adapt and refine the model for improved accuracy and performance. The primary challenge in linking ADM1da with metaproteomic data is that while metaproteome data can be mapped to MAGs, assigning these MAGs to specific metabolic pathways presents several difficulties: (i) The functional descriptions of microorganisms in the literature are often inferred from closely related species. These functions are typically determined in pure culture experiments or inferred solely from genomic data, which may not fully represent the metabolic roles of these organisms in complex environments like AD. (ii) Microorganisms potentially possess a wider range of metabolic functions, but their expression is heavily influenced by environmental conditions. As a result, many pathways may not be active or detectable under the specific conditions of AD. (iii) Assigning pathways is further complicated by the presence of similar genes in different microbial taxa. For instance, primary and secondary fermenters often share homologous genes, making it difficult to distinguish their specific contributions to metabolic processes. (iv) Certain microbial taxa specialize in breaking down monomers while leaving the hydrolysis of larger biomass molecules to other species contrasting with the concept of the ADM1 (Campanaro et al., 2020; Menzel et al., 2020).

## 2. Material & methods

Besides the ADM1da model, this work is based on laboratory data already published in our earlier study. All information on the laboratory data can be found in Heyer et al. (2024a). For this study, the process parameters of the plant (Figure 2) and the metagenome and metaproteome data were used.

**Figure 2:**
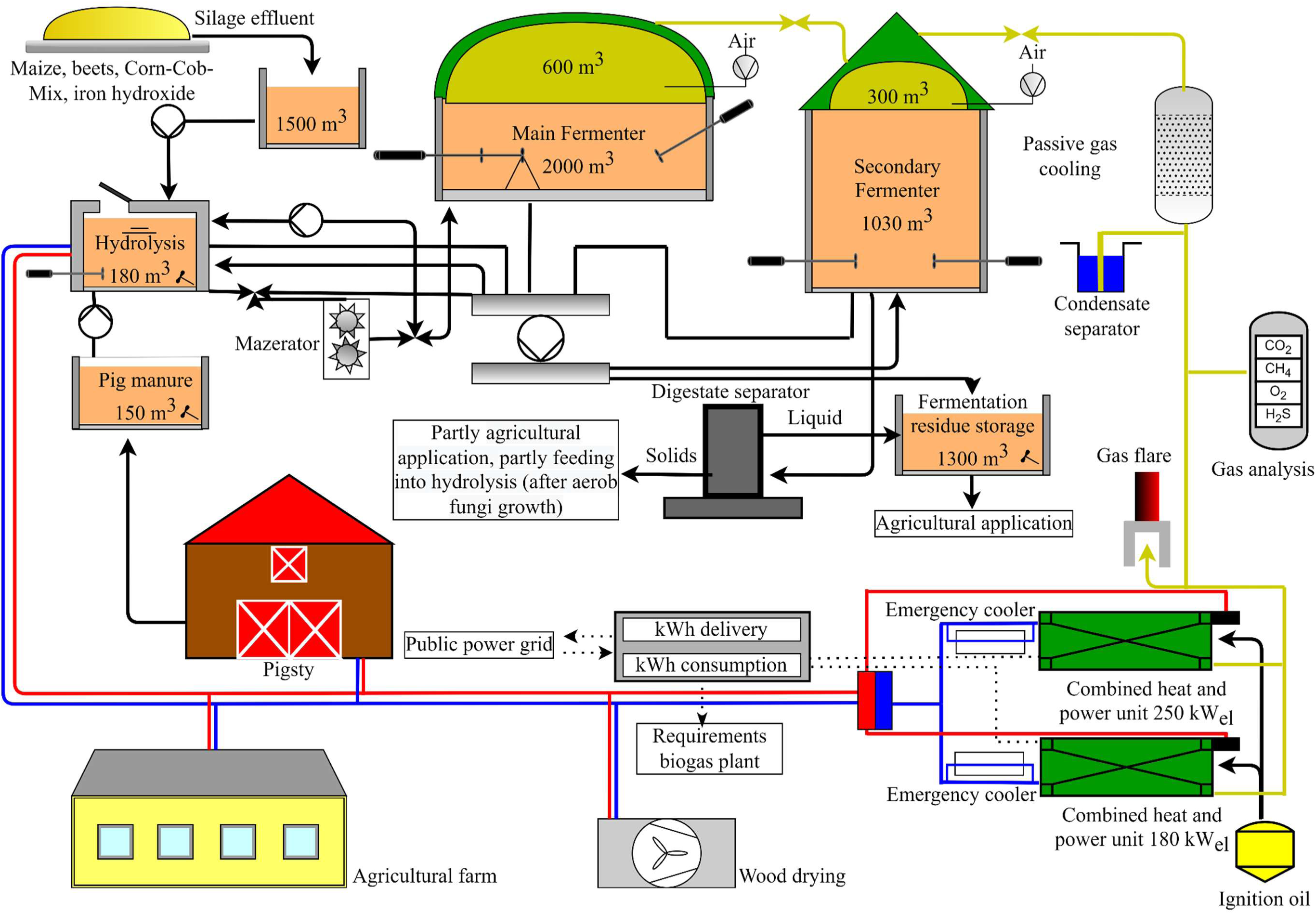
Schematic representation of the two-step anaerobic digestion plant. The system includes a hydrolysis reactor, a main reactor, and a secondary fermenter (not analyzed in this study). On average, 77.3 ± 5.0 tons of wet substrate were fed daily into the hydrolysis reactor, mixed with 49.2 ± 4.6 m^3^ of liquid fermentation residues from the secondary fermenter (see Supplementary Table 1). Blue and red lines indicate heat flow, dotted lines represent electrical current, black lines show the movement of substrates or residues, and yellow lines depict the production of biogas.

## Representation of the AD using Simba# and ADM1da

Simulations of the two-step AD were performed with the simulation software Simba# (Version 4.3) (Alex et al., 2013; ifak GmbH, 2021; Rojas et al., 2011) based on the ADM1da (Karlsson, 2017; Ogurek et al., 2013). Additionally, a single-step process was simulated in the absence of a hydrolysis fermenter to facilitate a comparative analysis of two-step processes (Figure 3).

**Figure 3:**
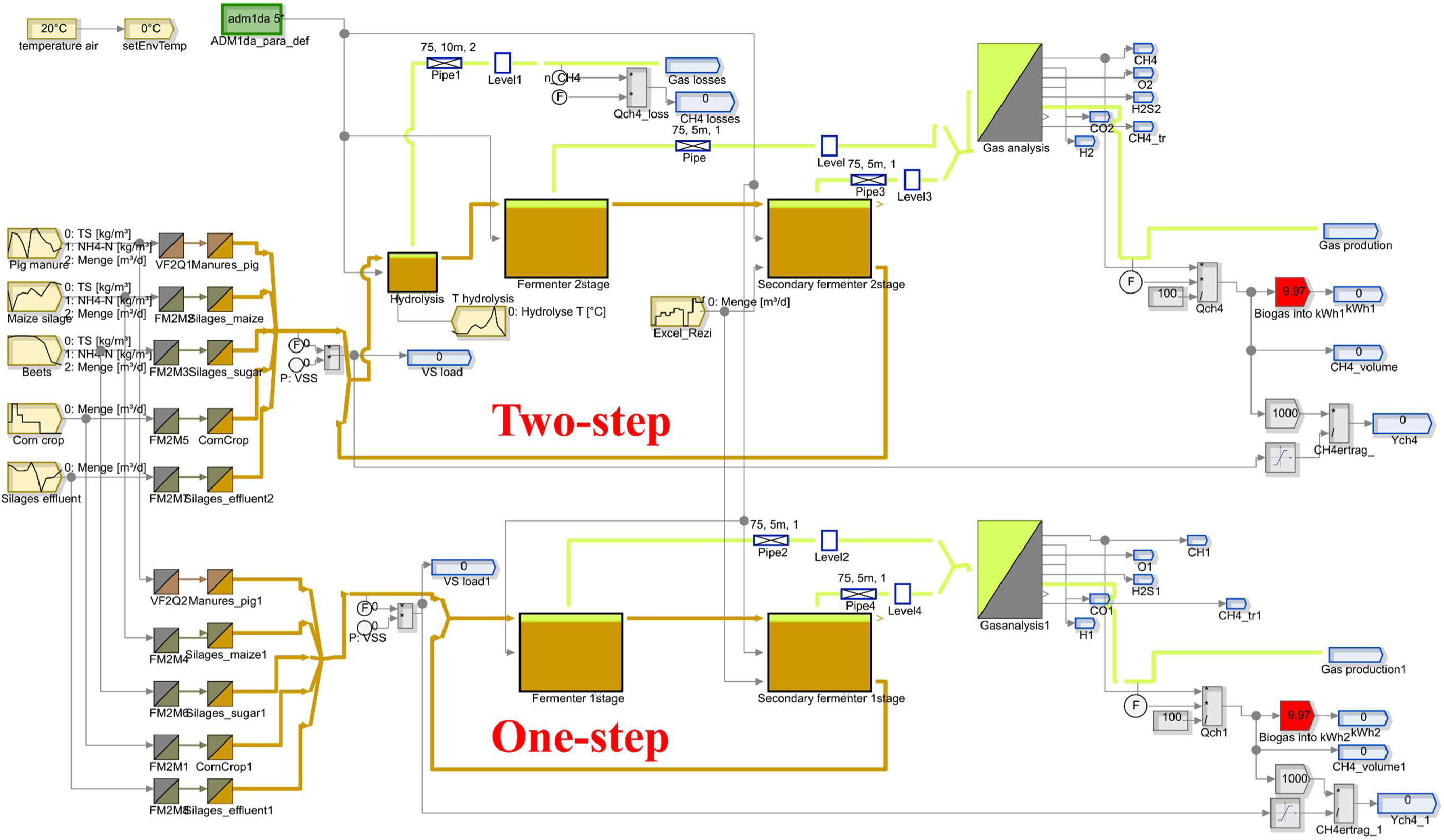
Structure of the Simba# model used for simulations: 1. Biomass input, 2. Converter block, 3. Hydrolysis fermenter, 4. Main fermenter, 5. Recirculation of biomass, 6. Secondary fermenter, 7. Gas analyzer, and 8. Calculations (methane amount, methane yield, energy content biogas).

The final SIMBA model (ifak GmbH, 2021) of the two-step AD comprised seven pre-implemented building blocks: the biomass input/substrates (1), the converter blocks to calculate the ADM1da fractions from the substrate data (2), the hydrolysis block (3), the main fermenter block (4), the secondary fermenter block (6) the recirculation block (5), the gas analyzer block (7) and blocks for calculation of methane amount, methane yield and energy content of biogas (8) (Figure 3). For the biomass input, the process parameters, and the AD design, we used the data provided by the plant operator (Supplementary Table 1), including the feeding, the temperature, and the fermenter sizes. Feeding information was forwarded to converter blocks of the SIMBA#, which are associated with a corresponding substrate model, the biomass input into a substrate composition utilized by the AMD1da (Supplementary Table 2).

For the input blocks of the simulation model, daily mean values of the measured feeding and recirculation amounts were used. The substrate qualities (total solids (TS) and volatile solids (VS)) were also given to the model as dynamic inputs according to the monthly laboratory measurements. For the final simulations, 365 days of lead time were simulated to adjust the system using constant settings (average values of feeding values), followed by 365 days of actual simulation matching to the actual analyzed period (described by daily values). To fit the simulation data to the actual AD data, the following three adaptations to the kinetic parameters of the SIMBA# implementations of ADM1da were made here - according to Seick et al. (2023): (i.) Increase of the upper limit for the modeled pH-limitation of methanogenesis (parameter pH_UL,ac_ in process p11 in ADM1da from 7.0 to 8.5); (ii.) an increase of the half-saturation constants for NH_3_ inhibition of propionic acid degradation and of acetoclastic methanogenesis (parameter K_INH3,pro_ in process p10 in ADM1da from 0.0019 to 0.0025 kmol_N_ m^-3^ and KI_NH3,ac_ in process p11 in ADM1da from 0.0018 to 0.0025 kmol_N_ m^-3^), (iii.) reduction of the acetoclastic methanogenic biomass decay rate (parameter k_dec,xac_ in process p18 in ADM1da from 0.04 to 0.02 d^-1^). These parameter changes are not a calibration for a specific case, but rather the application of suitable basic settings, which are recommended by Seick et al. (2023) as the basis for successful model adjustment.

As also described in Seick et al. (2023) the main part of the model adjustment was the setting of the inert fractions of VS of the substrates in the model so that the methane production of the plant was approximately achieved in the simulation. This inert fraction results indirectly from the parameterizations in the influent model by adjusting the parameters of biomethane potential (BMP), the crude fiber fraction of TS (fRF), and the degradable fraction of crude fiber (fOTSrf) in the substrate blocks of the respective substrates of the SIMBA# model (Figure 3).

## Assigning MAGs to the functional groups of the ADM1da

Annual abundances of microbial groups in the hydrolysis and the main fermenter were assessed by a metagenome-supported metaproteomics analysis covering twelve time points (Heyer et al., 2024a). The performed metaproteomics analysis quantified the relative abundances of microbial groups based on the unique spectral abundance of identified protein groups. Therefore, the spectral abundance of each protein group for each measurement was normalized to the total spectral abundance of the measurement. After grouping the MAGs into the seven groups defined by the ADM1da model (see below), we normalized the protein abundances of the MAGs within these groups to 100%. To maintain consistency in the biomass dynamic of the predicted COD values, the sum of all biomass COD averages was normalized over the entire period of one year. This ensures that the metaproteomics analysis and the COD biomass are comparable. The protein assignment to the microbial groups from the ADM1da was done based on the linkage of the proteins resp. genes to high-abundant and high-quality MAGs (>90% completeness, <5% contamination) (Heyer et al., 2024a) and the assignment of the MAGs to the seven microbial groups (Figure 1) based on the following rules. The first rule groups the MAGs according to the ADM1da groups based on the metabolic functions from the literature associated with each MAG’s taxonomy. MAGs with functions outside the seven groups of the ADM1da or with low taxonomic resolution (below the family level) were excluded. This rule is referred to as the “Literature Rule“ in the following. The second rule groups the MAGs based on the identified proteins associated with a pathway of the ADM1da, without consideration of taxonomy. This rule is referred to as the “Pathway Presence Rule“ in the following. The third rule called “Extended Pathway Rule“ builds on the “Pathway Presence Rule“, incorporating specific thresholds and taxonomic information to categorize the MAGs within ADM1da groups. The final rule, termed the "Fine-Tuned Pathway Rule," represents an enhancement of the "Extended Pathway Rule." It incorporates additional ratios and protein abundance thresholds, thereby further refining the grouping. This is done to distinguish between sugar and amino acid-degrading bacteria and between primary and secondary fermenting bacteria (Table 1 and Supplementary Table 3).

**Table 1:**
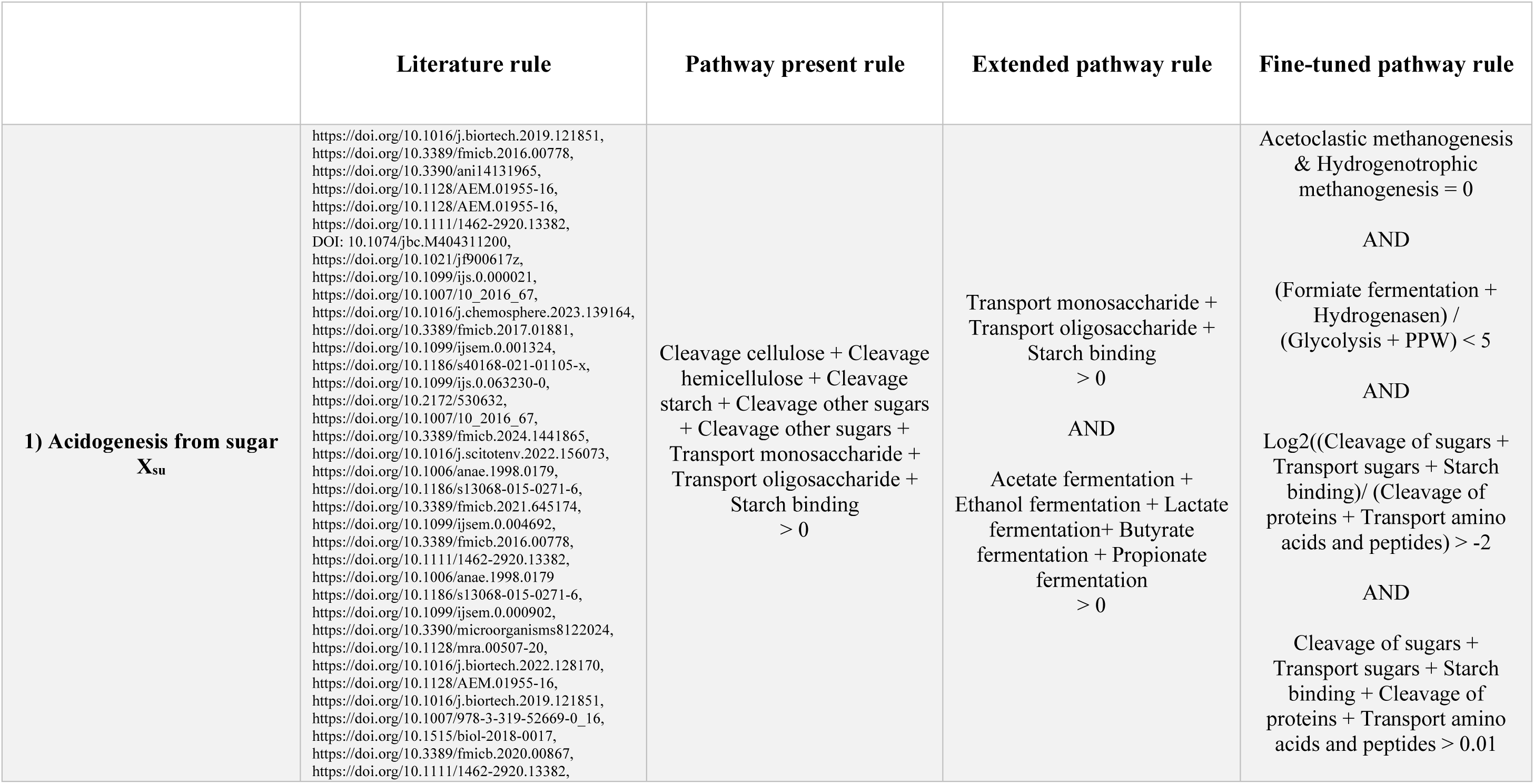
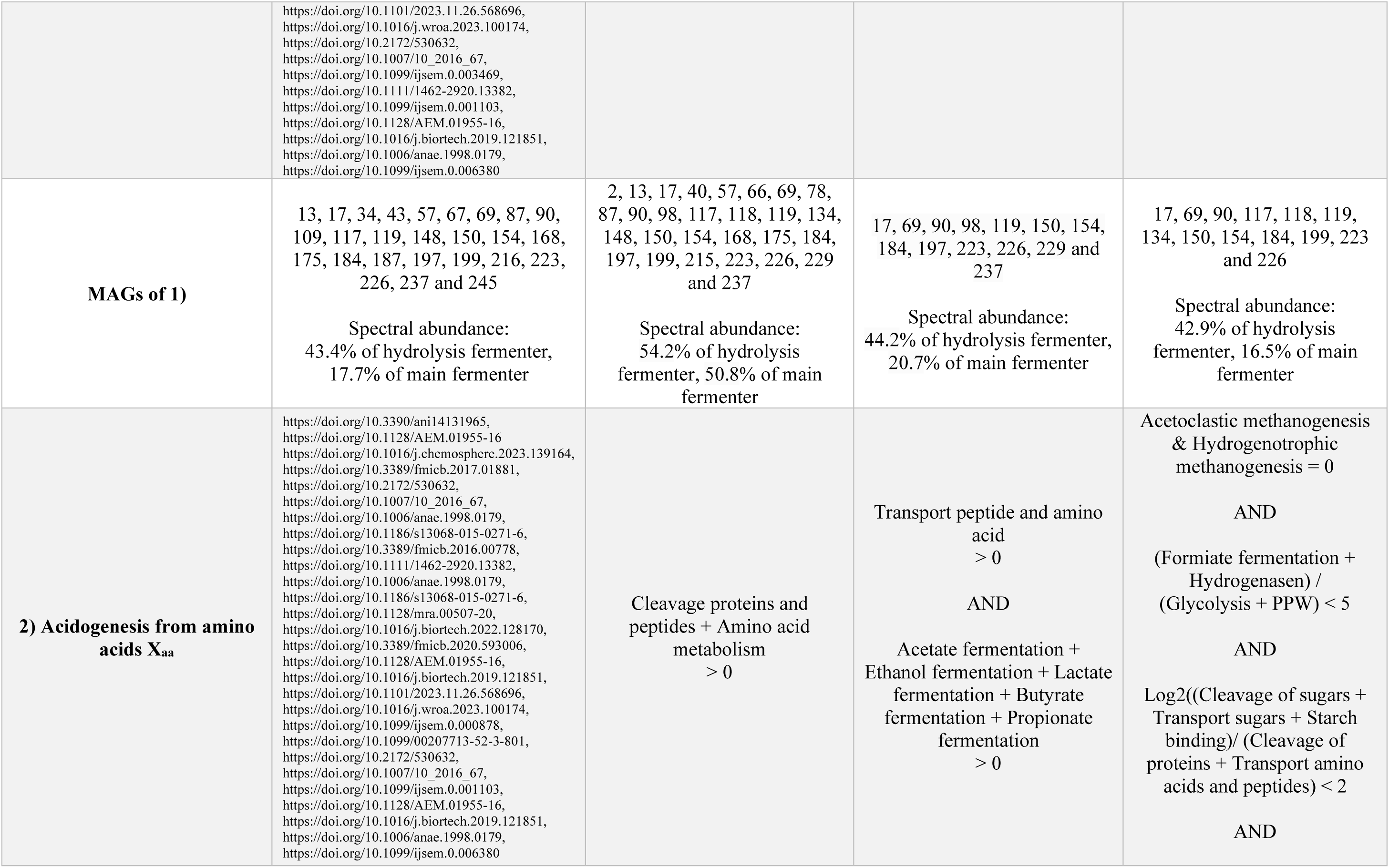
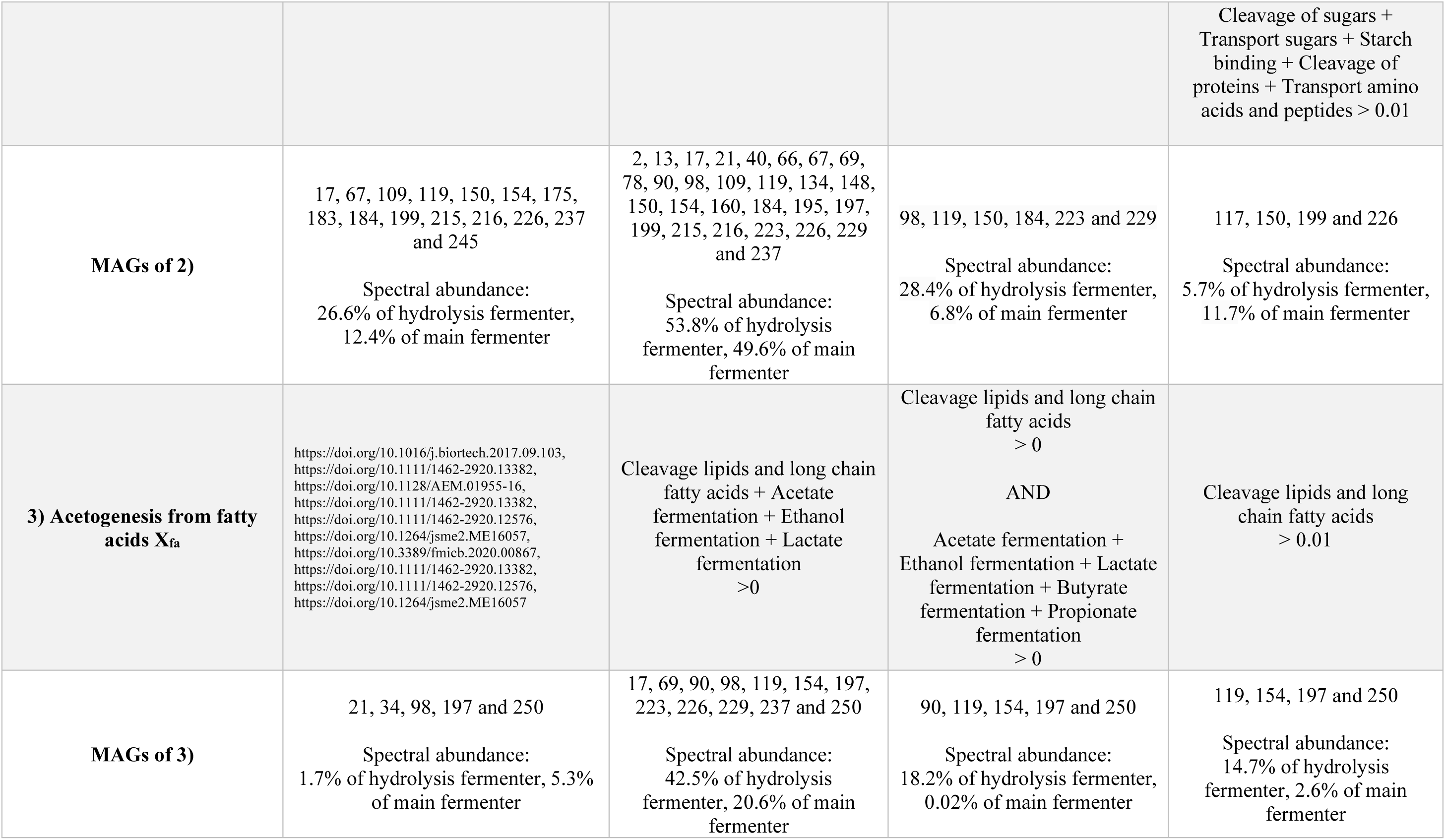
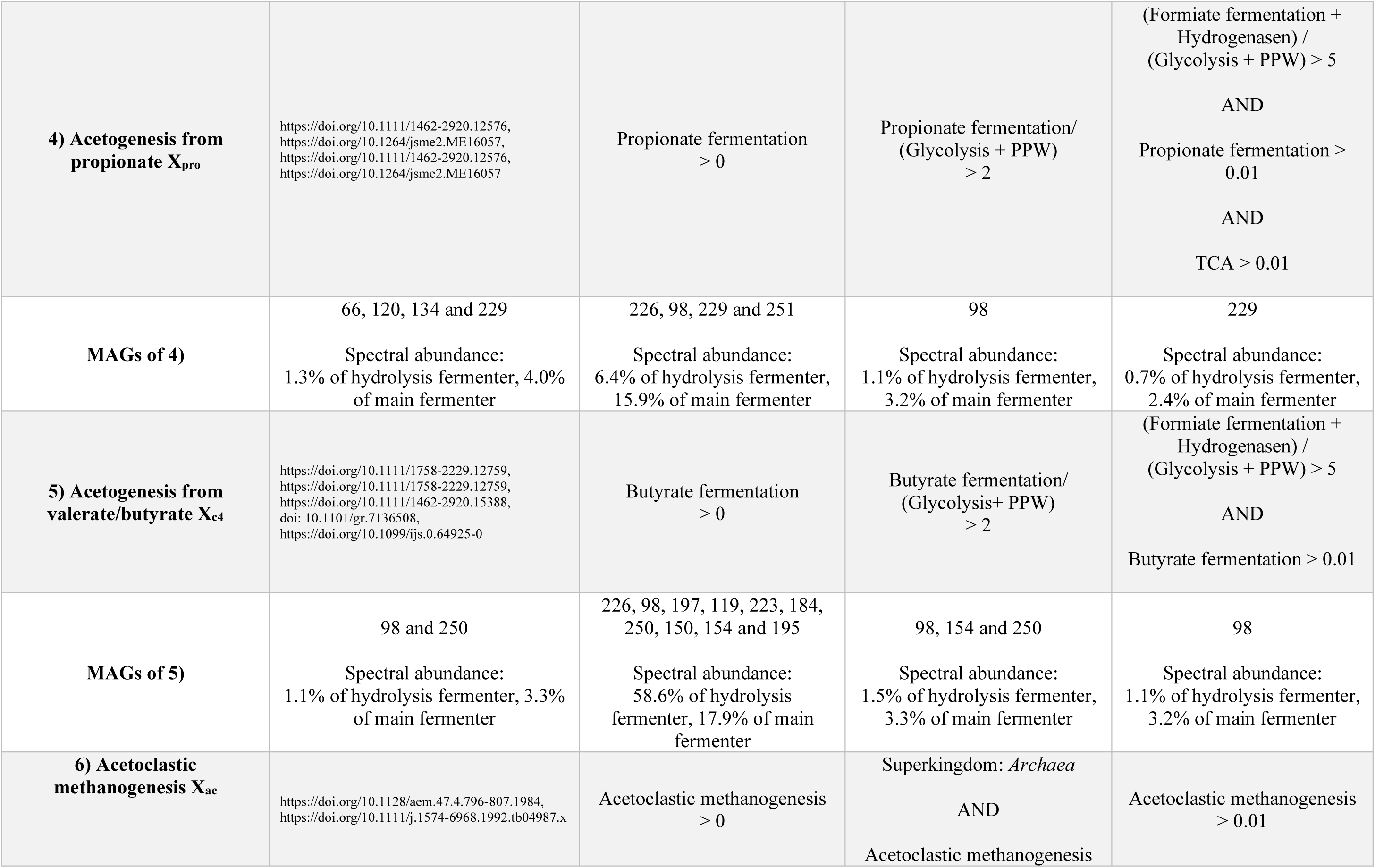
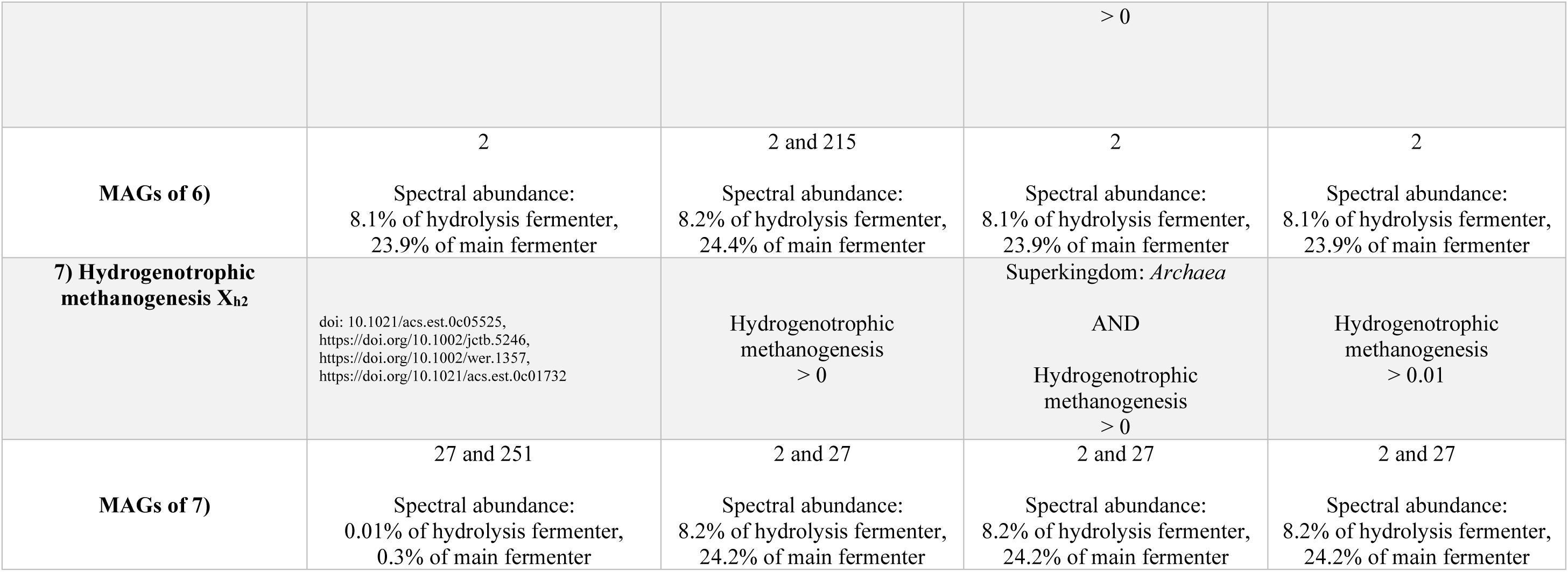
High-quality MAG assignment to the seven functional groups of the ADM1da based on protein data. MAGs were grouped based on four different rules. MAGs that fit into multiple groups based on their identified proteins were assigned to all relevant groups by dividing their abundance equally. The raw data can be found in Supplementary Table 4, PPW: pentose-phosphate pathway.

## 3. Results and Discussion

### Simulations using the ADM1da model of the two-step anaerobic digestion plant

The sampled two-stage anaerobic digester system is comprised of a non-gas-tight continuous stirred tank reactor for hydrolysis with a volume of 180 m³, and a main fermenter with a volume of 2000 m³ (Figure 2). On average, 77.3 ± 5.0 tons of wet substrate were fed daily into the hydrolysis reactor, mixed with 49.2 ± 4.6 m³ of liquid fermentation residues from the secondary fermenter (not analyzed in this study). The distinctive aspect of this plant is the incorporation of a hydrolysis fermenter that is designed to pre-mix the substrate and facilitate the pre-digestion and breakdown of complex substrates for the main fermenter. This represents a unique approach to AD. The process data about the hydrolysis and the main fermenter were employed as input for the ADM1da model (Figure 3).

To assess the model accuracy, we evaluated whether the exact representation of the ADM1da fitted the actual produced amount of biogas and methane (Supplementary Table 4). The average simulated produced amount of biogas by the model was 3,272.3 m^3^ day^-1^ ± 202.7 m^3^ day^-1^, compared to the measured abundance of 3,175.7 m^3^ day^-1^ ± 301.7 m^3^ day^-1^ (Figure 4 A), showing a good accordance. Similar predicted methane content in biogas was 56% ± 0.6% compared to the measured 60.2% ± 1.5% (Figure 4 B), and the modeled produced methane amount was 1,728.1 m^3^ day^-1^ ± 104.7 m^3^ day^-1^, compared to the measured abundance of 1,911.5 m^3^ day^-1^ ± 178.3 m^3^ day^-1^ (Supplementary Figure 1 A). Following this, the mean percentage error (MPE) and root-mean-square error (RMSE) (Hyndman and Koehler, 2006) were 3.6% and 7.9% for biogas production, -6.9% and 7.5% for methane content, and -9.0% and 12.4% for methane production (Supplementary Table 4).

**Figure 4:**
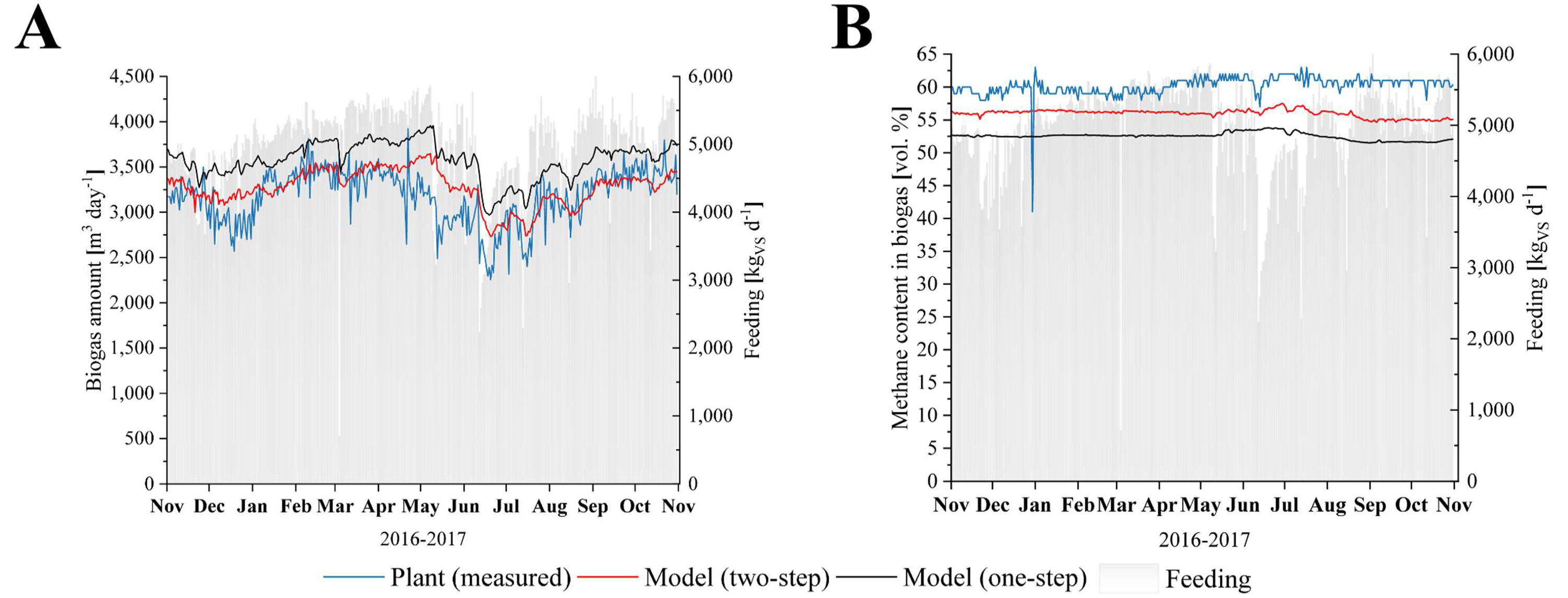
Comparison of the biogas amount and the methane content of the actual anaerobic digester compared to the ADM1da model. The simulation is based on daily values for feed quantities and monthly measured values for TS content or average values to describe the qualities of the different substrates. **A:** Simulation of the amount of biogas produced from the two-step AD with an open hydrolysis fermenter and a one-step AD where the hydrolysis fermenter is excluded. **B:** Simulation of the methane content produced from the two-step AD with an open hydrolysis fermenter and a one-step AD where the hydrolysis fermenter is excluded (see Supplementary Tables 2 and 4)

Furthermore, the monthly laboratory analyses of the concentrations of TS, VS, and volatile fatty acids (VFA), pH values, and the ammonium-nitrogen concentration (NH4-N, only available for the main fermenter) were reproduced sufficiently well by the model in the operating year under consideration (Supplementary Table 2, Supplementary Figure 2 and 3). In particular, the simulated VS degradation can serve as a plausibility check for the correct simulation of gas production. The comparison of the VS concentrations measured in the fermenters (VS_H_: 81.72 kg m^-3^ ± 9.00 kg m^-3^, VS_F_: 42.45 kg m^-3^ ± 5.00 kg m^-3^) with the monthly monitoring with the simulated time course (VS_modelH_: 88.77 kg m^-3^ ± 5.37 kg m^-3^, VS_modelF_: 41.73 kg m^-3^ ± 3.81 kg m^-3^) shows a good agreement.

Despite the sufficient fit of the ADM1da, there are still deviations in some parameters. The deviations between the simulated and measured values for pH (-13.87%) and VFA (14.92%) in the hydrolysis tank (Supplementary Figure 2) in the first half of the investigation period may be due to an initial state of the simulation that has not yet been reproduced with sufficient accuracy. Solving this issue would require additional operational data before the investigation period (at least for several months), which was not possible within the scope of the study. In addition, the model may not have fully captured the effects of substrate changes (e.g. stopping sugar beet feeding from April) on the process biology. Further reasons for deviations from simulation and measurement - especially in the methane content of the biogas and the methane production derived from it - are measurement inaccuracies in online gas analysis. A plausibility check (involving the summation of the gas concentrations measured online) revealed a measurement error of approximately 5% (due to a shift in April) by the end of the investigation period. This measurement error would correspond to an approximate 2-3% increase in the measured values for the CH_4_ content (Supplementary Figure 4 and Supplementary Table 1). The model may also underestimate CO_2_ outgassing in the open hydrolysis fermenter, which would further explain the lower simulated CH_4_ content in the biogas.

## Consequences of open hydrolysis and comparison with a single-step system

The model predicted a release of 430.9 m^3^ day^-1^ ± 84.5 m^3^ day^-1^ biogas and 45.8 m^3^ day^-1^ ± 33.6 m^3^ day^-1^ (10.6%) methane in the open hydrolysis fermenter, which was lost due to the missing gas collection in the real AD. The methane content and thus the methane losses increased significantly in the simulation with the establishment of the methanogens in the hydrolysis fermenter, especially from April onwards (see Supplementary Figure 1 B). If these methane losses are related to the simulated methane amount from the fermenter and secondary digester (1,728.1 m^3^ day^-1^), an average methane loss of 2.6% is obtained via the open hydrolysis fermenter. Matching this, the original paper identified larger amounts of enzymes for methanogenesis in the hydrolysis fermenter (Heyer et al., 2024a). Since the methane loss of about 45.8 m^3^ day^-1^ in the open hydrolysis is associated with economic losses and the promotion of global warming, the concept of an open hydrolysis fermenter at pH values around 5.9 is not promising.

Due to the methane loss in the open hydrolysis, ADs’ methane production was reduced (Figure 4 and Supplementary Figure 1). However, in the case the biogas from the hydrolysis fermenter is collected, the biogas production would increase from 3,272,3 m^3^ day^-1^ to 3,703,2 m^3^ day^-1^ (+13.2%), and the methane production from 1,728.1 m^3^ day^-1^ to 1,773.8 m^3^ day^-1^ (+2.6%). Simulations of the same AD without the hydrolysis fermenter revealed a biogas production of 3,575.7 m^3^ day^-1^ (-3.6% compared to the two-step AD with gas-tight hydrolysis fermenter) and a methane production of 1,771.9 m^3^ day^-1^ (-0.1%). This indicates that a two-step system with closed hydrolysis would be only slightly more effective than a single-step system of the same volume.

## Linking metaproteomics data to the ADM1da

Metaproteomics data were linked to MAGs and subsequently assigned to functional groups based on MAGs. Due to challenges in linking the data, we developed four rules: (i) “Literature Rule“, (ii) “Pathway Presence Rule“, (iii) “Extended Pathway Rule“, and (iv) “Fine-Tuned Pathway Rule“ (Table 1). To assess the quality of the metaproteomics assignment to the ADM1da model, we compared the number of linked high-abundance MAGs and evaluated the MPE between the model and data. This approach allowed us to estimate the accuracy of the linkage. Additionally, we used the Spearman correlation between the model and the measured biomass to assess how well the model dynamics fit the data.

The assignment of MAGs and their associated spectral abundance to the seven microbial groups in the ADM1 model enabled the linkage of 38 MAGs based on (i) “Literature Rule“, 37 MAGs based on (ii) “Pathway Presence Rule“, 16 MAGs based on (iii) “Extended Pathway Rule“, and 19 MAGs based on (iv) “Fine-Tuned Pathway Rule“ (Table 1). These assignments were derived from 49 high-quality MAGs (completeness > 90%, contamination < 5%) according to Bowers et al. (2017). The 49 high-quality MAGs represented 52.7% of the identified spectra in the main fermenter and 54.8% in the hydrolysis fermenter. The 49 high- quality MAGs accounted for 52.7% of the identified spectra in the main fermenter and 54.8% in the hydrolysis fermenter. Notably, 9 of these MAGs represented already 47.66% of the identified spectra in the main fermenter (abundance ≥ 1%), and 8 MAGs accounted for 48.95% in the hydrolysis fermenter (abundance ≥ 1%) (Table 1 and Supplementary Table 3). This indicates that 8-9 MAGs comprise approximately half of the biomass in this AD on average. The highest accuracy (median of MPE ± average absolute deviation of MPE) in terms of model abundance was achieved using the (ii) “Pathway Presence Rule“ (MPE for hydrolysis: 67.22% ± 41.71%, MPE for fermenter: 54.26% ± 28.73%), followed by the (iii) “Extended Pathway Rule“ (MPE for hydrolysis: 56.21% ± 77.35%, MPE for fermenter: 59.24% ± 23.39%). The (iv) “Fine-Tuned Pathway Rule“ (MPE for hydrolysis: 61.78% ± 35.93%, MPE for fermenter: 51.27% ± 96.99%) and (i) “Literature Rule“ (MPE for hydrolysis: 52.50% ± 1235.67%, MPE for fermenter: 50.46% ± 309.20%) showed lower accuracy (Figure 5 and Supplementary Table 5).

**Figure 5:**
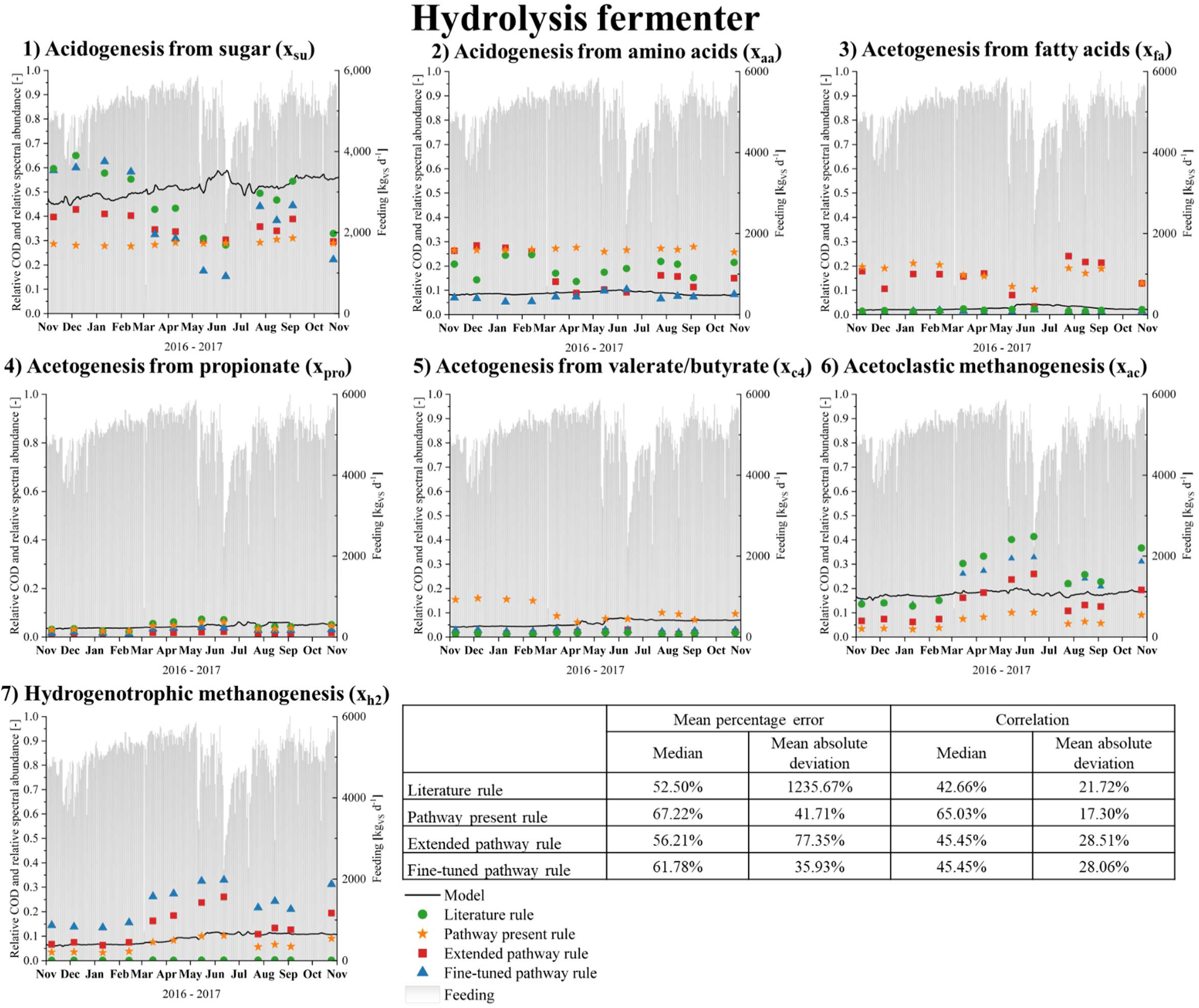
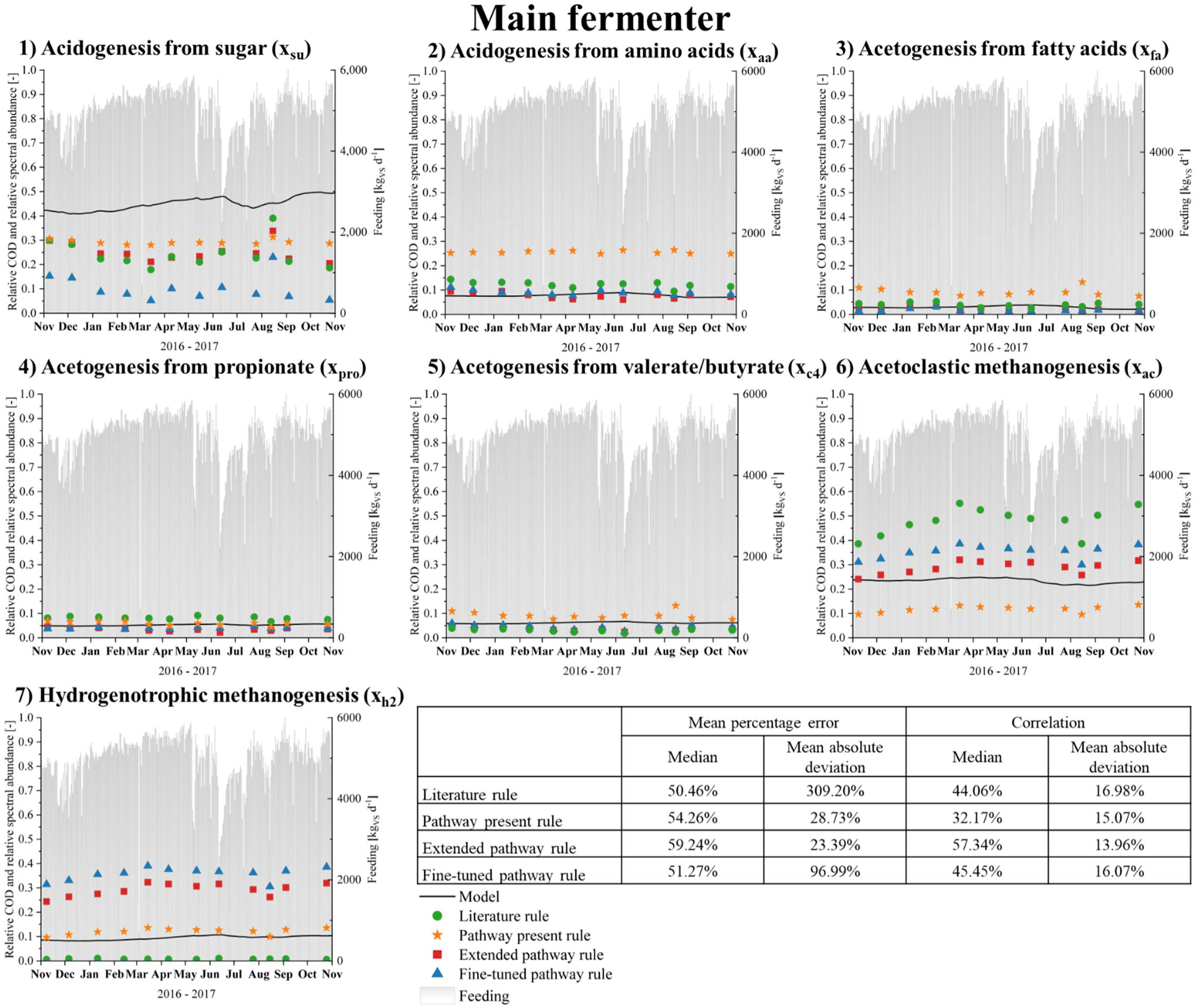
Comparison between predicted and measured biomass composition of the seven functional groups of the ADM1da model. Following the grouping of MAGs into the seven categories defined by the ADM1da model (see Table 1). To allow a direct comparison with metagenome abundance, protein abundances of the MAGs in each group were normalized to 100%. Similarly, the predicted COD values for ADM1da groups were normalized to 100%. Protein assignments to the microbial groups in ADM1da are based on the association of proteins or genes with high- abundance, high-quality MAGs (≥90% completeness and ≤5% contamination). The MPE was calculated based on residuals and the correlation was calculated using Spearman’s correlation. For clarity, only the median and mean absolute deviation of the MPE and correlation values across the various rules are shown in this figure. The MPE and correlation values for each microbial group in the ADM1 model, based on each rule applied, are shown in Supplementary Table 5. All further information can be found in Supplementary Tables 3 and 5.

Conversely, for the dynamic correlation (median of Spearman’s rank correlation coefficient ± average absolute deviation of Spearman’s rank correlation coefficient), the worst performance was observed for the (ii) “Pathway Presence Rule“ (correlation for hydrolysis: 65.03% ± 17.30%, correlation for fermenter: 32.17% ± 15.07%). The highest correlations were obtained with the (iii) “Extended Pathway Rule“ (correlation for hydrolysis: 45.45% ± 28.51%, correlation for fermenter: 57.34% ± 13.96%), followed by the (iv) “Fine-Tuned Pathway Rule“ (correlation for hydrolysis: 45.45% ± 28.06%, correlation for fermenter: 45.45% ± 16.07%) and (i) “Literature Rule“ (correlation for hydrolysis 42.66% ± 21.72%, correlation for fermenter: 44.06% ± 16.98%) (Figure 5 and Supplementary Table 5).

While the results are not yet optimal, we have successfully achieved an acceptable mapping of the ADM1da model to the microbial abundance data. For example, we were able to achieve a correlation of over 90% and an MPE of less than 10% in some groups. However, it is notable that some rules demonstrate a high degree of accuracy in representing certain groups while exhibiting a lower level of precision in representing other groups.

This mapping, along with a deeper analysis of MAGs, provides novel insights into actual metabolic processes. More sophisticated models could now be developed based on this functional analysis. However, a perfect mapping of the data to ADM1da cannot be expected due to several issues where the model fails to capture the actual microbiome dynamics:

(1) Abundance and dynamics of sugar, amino acid, and fatty acid degraders: We observed the greatest variance here. Several MAGs (e.g., MAG_98 family *Syntrophomonadaceae* and MAG_154 genus *Eubacterium_H*) are divided into one to four groups, depending on the rule. This uncertainty highlights that some microorganisms don’t fit neatly into a single group, leading to ambiguous classifications. We have evenly distributed the MAGs across groups, though it’s uncertain if this is the optimal approach but data is still missing to make the classification even more accurate. Furthermore, some MAGs involved in sugar degradation have been observed to express only proteins associated with monomer degradation (e.g., MAG_117 species *GCA_001512825.1* and MAG_229 species *GCA_002305915.1*), while a few (e.g., MAG_69 species *Herbinix luporum* and MAG_78 family *Chitinispirillaceae*) have been shown to encode enzymes essential for the hydrolysis of complex carbohydrates. Nevertheless, all these MAGs have been classified within the same model group for acidogenesis from sugar.

The model is based on the assumption of mostly stable conditions throughout the year; however, the protein data demonstrate notable fluctuations. Furthermore, the model does not account for changes in substrate composition, such as the cessation of sugar beet feeding in April, which has a marked impact on the protein abundance of specific MAGs (e.g., MAG_229 and 223). Changes in feedstock composition (e.g., sugar and amino acids) lead to microbial shifts that are not captured by the model. Protein data offer invaluable insights into these overlooked microbial and metabolic changes, as demonstrated in our study. These unrecorded dynamics exerted a considerable influence on the MPE and the correlation coefficients.

(2) Assignment issues in acetoclastic and hydrogenotrophic methanogenesis: As documented in the literature, *Methanotrix* (MAG_2), which constitutes 24% of the biomass and is the most abundant MAG of this AD, is a strictly acetoclastic *archaea*. However, the expressed proteins suggest that *Methanotrix* may also perform hydrogenotrophic methanogenesis (Heyer et al., 2024a; Khesali Aghtaei et al., 2022). Consequently, its group assignment exerts a significant influence on the observed fluctuations. For example, in “Literature rule”, this MAG causes by far the greatest error, because of the strict assignment to the acetoclastic group (Figure 4). Furthermore, the presence of cytochromes influences energy yield and biomass formation in both acetoclastic (1.5 mol ATP with cytochrome per 1 mol acetate vs. 1 mol ATP without) and hydrogenotrophic methanogens (1.5 mol ATP with cytochrome per 1 mol CH_4_ produced vs. 0.5 mol ATP without) (Jin, 2012; Thauer et al., 2008). In the future, this may be recorded using omics data, thereby facilitating a more accurate prediction of archaeal activity (Basile et al., 2023).

(3) Unlinked abundant MAGs: Despite successfully linking key species representing AD to ADM1da, some highly abundant MAGs, such as MAG_78 (1.77%), MAG_113 (2.97%), MAG_117 (1.29%), and MAG_153 (1.82%), were not linked to the ADM1da model. MAG_113 may act as a fermenting bacterium, but its sequence quality is low, as is MAG_153, which may be a giant virus (A32-like ATPase (A32)) (Mavian et al., 2012). MAG_78 and MAG_117 are high-quality MAGs identified as syntrophic acetate-oxidizing (SAO) bacteria, organisms not included in ADM1da (Basile et al., 2023; Capson-Tojo et al., 2021). However, SAO bacteria are indispensable for biogas production, particularly in mitigating the inhibitory effects of elevated ammonia levels on methanogenesis (Westerholm et al., 2012). Furthermore, studies show that ADM1da lacks pathways for lactate and alcohol fermentation, which limits its accuracy in certain contexts (Heyer et al., 2019; Shi et al., 2019). It has been observed that high organic loading rates and high-sugar substrates, such as food waste, can promote increased lactate and alcohol formation (García-Depraect et al., 2022; Moestedt et al., 2019). These further highlight potential areas where ADM1da may benefit from refinement based on proteome analysis.

(4) Limited resolution of metagenomics and metaproteomics: Current sequencing and proteomics methods still restrict our ability to fully decipher microbiome complexity. For instance, only 50% of the biomass was comprised of high-quality MAGs, while 25% of the biomass could not be assigned to any MAG. However, this limitation is expected to decrease as sequencing depth improves (Hu et al., 2021), along with the advent of novel mass spectrometry techniques and DIA-based metaproteomics workflows (Dumas et al., 2024; Gómez-Varela et al., 2023). Matching to this, it was only feasible to calculate the abundance of microbial groups based on the spectral abundance of all proteins in a MAG, rather than focusing on marker proteins like ribosomal proteins (Suh et al., 2005), which could potentially offer better quantification (Starke et al., 2022).

## Future research at the interface between modelling and metaproteomics

Our study demonstrates that the ADM1da model effectively captures the key microbial community structure within the main fermenter, though several discrepancies arise due to processes not currently integrated into ADM1da. For instance, alternative metabolic pathways such as ethanol and lactate fermentation, as well as SAO, were shown by protein data. Additionally, certain microbial species participate in multiple functional roles, adding complexity to their representation. Despite these limitations, the correlation between metaproteomic data and ADM1da highlights the predictive potential of metaproteomics in elucidating ongoing metabolic processes. As mass spectrometry and bioinformatics continue to advance, enhanced metaproteomic resolution will provide deeper insights into anaerobic digestion (AD) stages. This could extend to rate-limiting processes such as hydrolysis limiting (Basile et al., 2023; Mo et al., 2023; Wang and Li, 2014), where the abundance of hydrolytic enzymes remains near the detection limit of an MS, leading to significant data variability and hindering their inclusion in this study.

Further refinement of ADM1da would benefit from examining metabolic fluxes of various microbial populations to clarify their specific metabolic roles and interactions (Basile et al., 2023; Bernardini et al., 2022; Koch et al., 2016; Lange et al., 2024; Weinrich et al., 2019). This integration would leverage metaproteomics’ ability to link functional and phylogenetic data, unveiling key enzymes and pathways involved in converting biomass to methane (Heyer et al., 2015). Such data could also simplify ADM1da by reducing reliance on parameters such as yield coefficients, thereby enhancing both its applicability and accuracy (Weinrich et al., 2019).

Our data suggest that substrate shifts can lead to changes in the microbial community that were not captured by the current model. Integrating these shifts with operational parameters and including additional groups, such as SAO, could enhance the model’s predictive power—not only for key process parameters but also for identifying potential process disruptions that might threaten microbial viability. These disruptions extend beyond established factors, such as high ammonium levels or low pH, to include less understood influences like microbial mortality rates. This mortality is driven by interactions with bacteriophages (Rossi et al., 2022), antimicrobial peptides (Wolak et al., 2023), and antibiotics (Visca et al., 2021), all of which could be quantified through metaproteomic approaches, offering new insights into community stability and resilience (Heyer et al., 2024b).

## 4. Conclusions

The ADM1da model demonstrates considerable effectiveness in simulating biogas and methane production within two-step AD processes, showcasing its potential alongside its limitations. Two-step AD systems, particularly those incorporating a hydrolysis fermenter, achieve slightly higher methane yields compared to one-step systems. However, these benefits are counterbalanced by methane losses from open hydrolysis fermenters, emphasizing the importance of employing gas-tight designs to minimize emissions. The integration of high-resolution datasets with advanced modeling frameworks is crucial for optimizing AD processes and promoting innovation in this field. Incorporating metaproteomics data into the ADM1da model offered valuable insights into microbial roles, yet the approach was constrained by unlinked MAGs and the omission of critical metabolic pathways, such as SAO, lactate fermentation, and alcohol fermentation. This study underscores the need to refine ADM1da by integrating these missing pathways and accounting for dynamic microbial community shifts, thereby enhancing the model’s accuracy and expanding its applicability, including the potential prediction of process disturbances.

## 6. Supplementary

Supplementary Note

Supplementary Table 1: All process parameters

Supplementary Table 2: Model fit process parameters

Supplementary Table 3: Rules for grouping MAGs into the ADM1da model

Supplementary Table 4: Model fit biogas values

Supplementary Table 5: Derivations between the ADM1da model and MAGs

## 8. Funding

This work was supported by the German Federal Ministry of Food and Agriculture (grants nos.22404015 and 22404115 and the German Federal Ministry of Education and Research (de.NBI network. project MicroGenUniBi. Grant no. 031L0103). We acknowledge support by the Open Access Publication Fund of Magdeburg University.

## 9. Data Availability Statement

Data Availability Statement: Proteome data were stored on PRIDE with the accession number: PXD044302 (http://www.ebi.ac.uk/pride/archive/projects/PXD044302)

Metagenome datasets were stored on Ena under the project number PRJEB39821. The reads were stored under the number accessions ERS15898928 through ERS15898933. The assembled metagenome data were stored under the number accession ERS17165236.

## 10. Conflicts of Interest

Not applicable.

## 11. Author contributions

- Conceptualization: R.H., P.H., I.S.
- Project administration: R.H., D.B., U.R.
- Sampling and characterization and process parameters: P.H.
- Experiments metaproteomics: P.H.
- Data evaluation: P.H., R.H., I.S.
- Bioinformatics: P.H., R.H.
- ADM1 modeling; I.S., N.M.
- Supervision: D.B., J.W., U.R.
- writing—original draft: P.H.
- writing—review and editing: I.S., R.H., D.B., J.W., U.R.

All authors have read and agreed to the published version of the manuscript.

## 12. Declaration of generative AI in scientific writing

ChatGPT and Grammarly were used to improve spelling and expression.

## Abbreviations

AD: Anaerobic digester, anaerobic digestion
ADM1: Anaerobic Digestion Model Number 1
ADM1da: Anaerobic digestion model number one da (Extension of ADM1)
BMP: Biomethane potential
COD: Chemical oxygen demand
fOTSrf: Degradable fraction of crude fiber
fRF: Fiber fraction of total solids
MAG: Metagenome-assembled genome
MPE: Mean percentage error
NH4-N: Ammonium-nitrogen concentration
ODE: Ordinary differential equation
OLR: Organic loading rate
rRMSE: Root-mean-square error
SAO: Syntrophic acetate oxidizing
VFA: Volatile fatty acids
TS: Total solids
VS: Volatile solids

## Notes

### Competing Interest Statement

The authors have declared no competing interest.

