## Supplementary Note for "Creating microbiome-model harmony between metaproteomics data and the ADM1da for a two-step anaerobic digester"

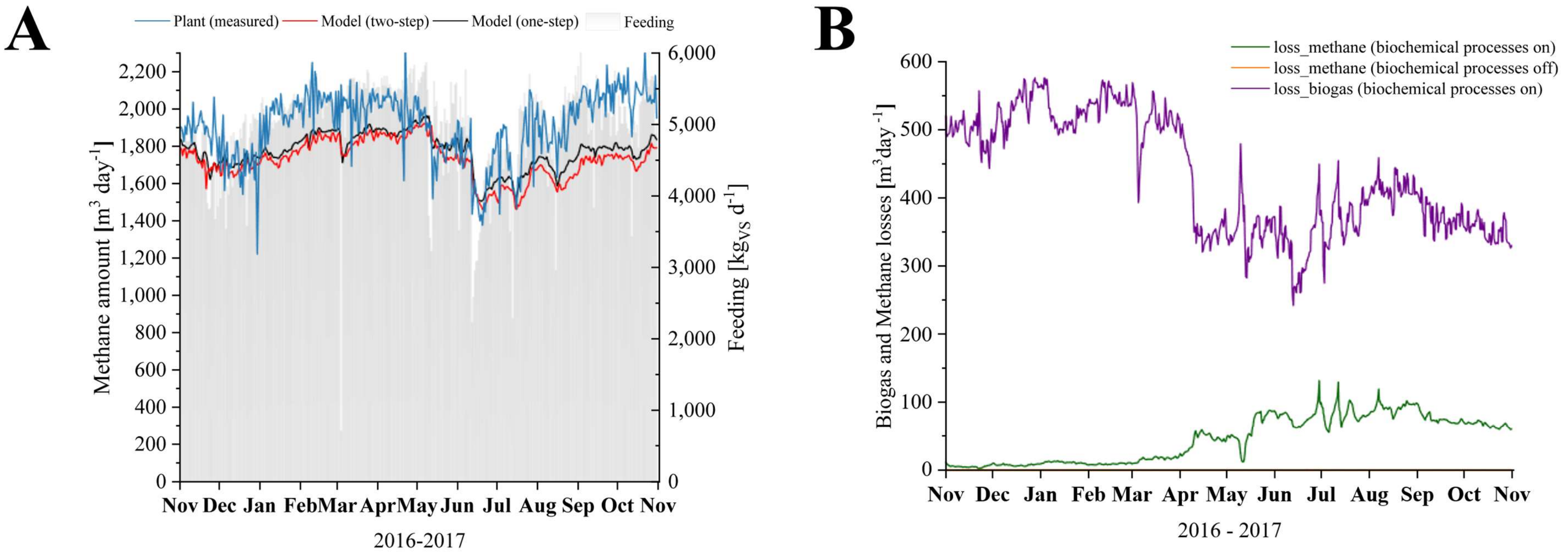


Supplementary Figure 1: Comparison of the methane output of the actual anaerobic digestion compared to the ADM1 model and the assumed gas losses of the ADM1 model. The simulation is based on daily values for feed quantities and monthly measured values for total solid contents or average values to describe the qualities of the different substrates (see Supplementary Table 1). **A:** Simulation of the amount of methane produced from the two-step AD with an open hydrolysis fermenter and a one-step AD where the hydrolysis fermenter is excluded. **B:** The estimated losses of the two-step AD due to the open hydrolysis fermenter. For the losses, the biochemical processes were turned on and off. The losses with biochemical processes were between 0 and 0.12. Therefore, it was excluded from the graph (see Supplementary Table 4).

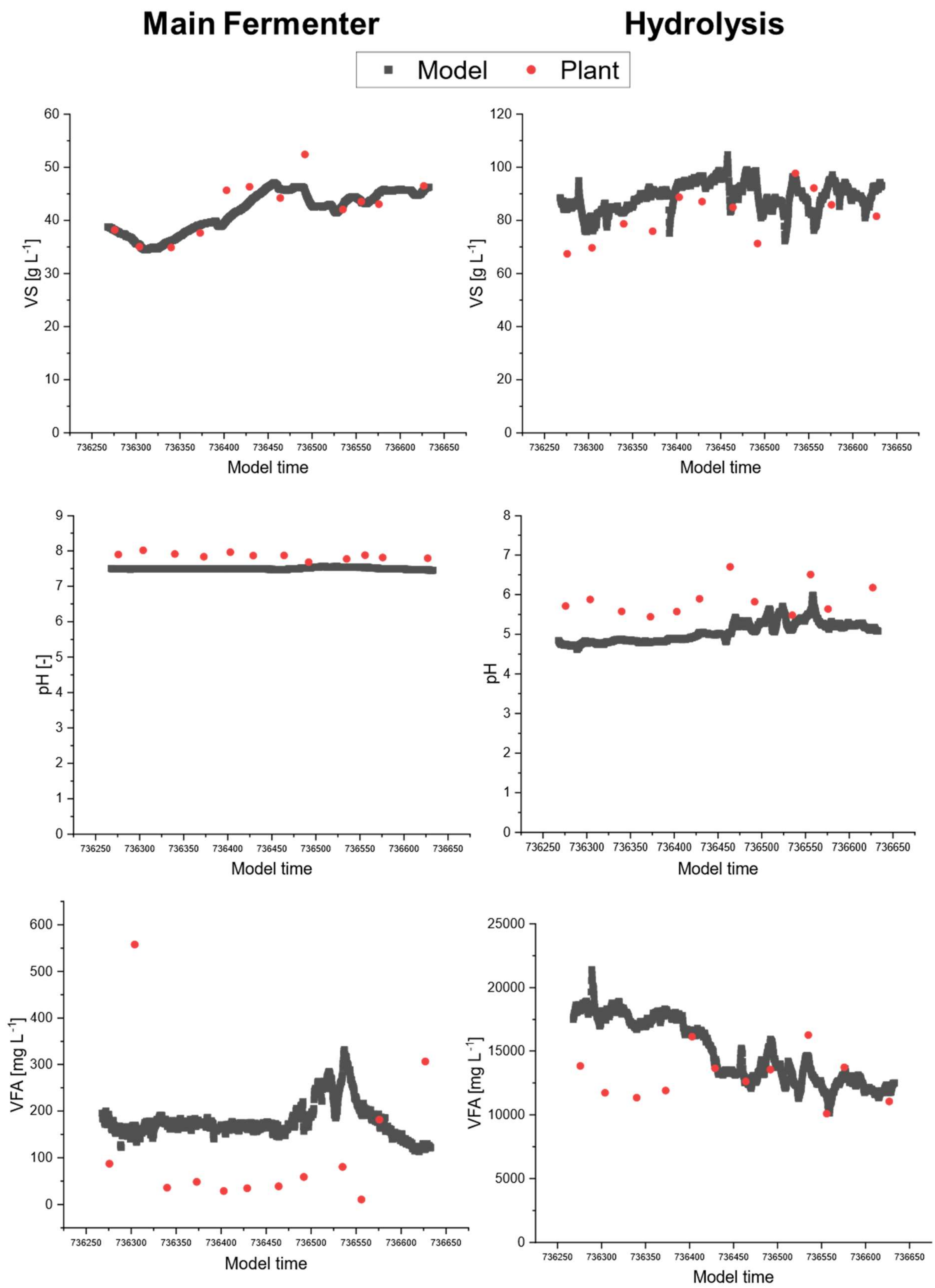


Supplementary Figure 2: Adaptation of the model to the chemical parameters VS, pH, and VFA of the plant. The daily values of the model were compared with the monthly averages of the plant data. Raw data can be found in Supplementary Table 2.

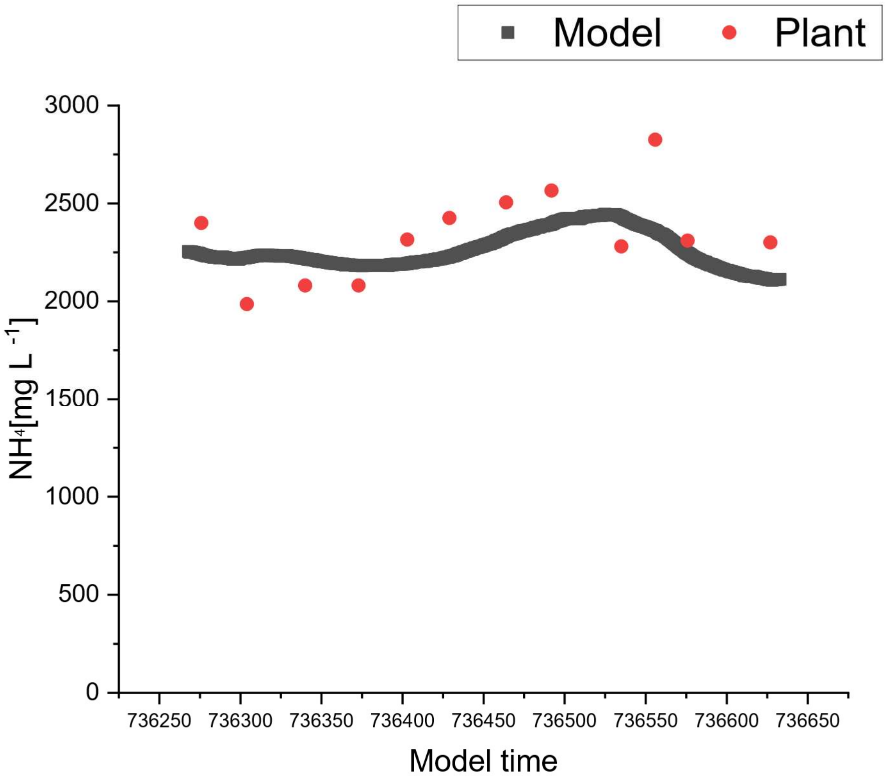


Supplementary Figure 3: Adaptation of the model to the ammonia-nitrogen content of the plant. The daily values of the model were compared with the monthly averages of the plant data. Raw data can be found in Supplementary Table 2.

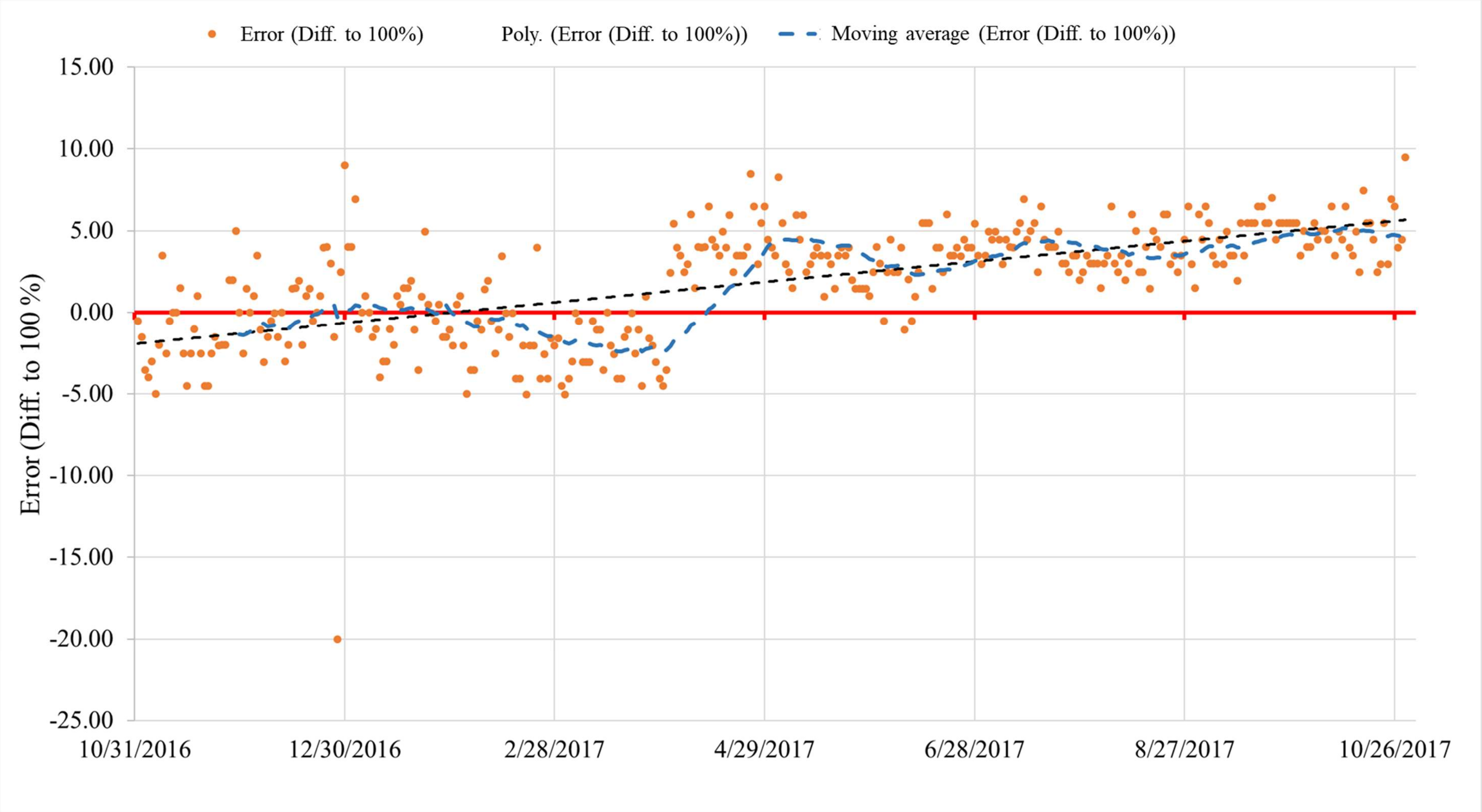


Supplementary Figure 4:Plausibility check of the performed gas analysis from the operator of the plant. Based on the volume percentages of the gas, the deviation was calculated to 100% (error to 100%). Based on the measurement error, the moving average and a polynomial for mapping the error curve were calculated. See also Supplementary Table 1.
